# MSRCall: A Multi-scale Deep Neural Network to Basecall Oxford Nanopore Sequences

**DOI:** 10.1101/2021.12.20.471615

**Authors:** Yang-Ming Yeh, Yi-Chang Lu

## Abstract

MinION, a third-generation sequencer from Oxford Nanopore Technologies, is a portable device that can provide long nucleotide read data in real-time. It primarily aims to deduce the makeup of nucleotide sequences from the ionic current signals generated when passing DNA/RNA fragments through nanopores charged with a voltage difference. To determine the nucleotides from the measured signals, a translation process known as basecalling is required. However, compared to NGS basecallers, the calling accuracy of MinION still needs to be improved.

In this work, a simple but powerful neural network architecture called MSRCall is proposed. MSRCall comprises a multi-scale structure, recurrent layers, a fusion block, and a CTC decoder. To better identify both short-range and long-range dependencies, the recurrent layer is redesigned to capture various time-scale features with a multi-scale structure. The results show that MSRCall outperforms other basecallers in terms of both read and consensus accuracies.

## 1 Introduction

In 2014, Oxford Nanopore Technologies (ONT) proposed a brand new nanopore sequencing product called MinION, featuring high-throughput single-molecule long read sequencing on a portable device. Compared to the next-generation sequencing (NGS) technology, MinION can produce sequences of lengths up to approximately 100 kbp, while most NGS methods only provide sequences up to several hundred bases long. The core of MinION is a membrane with up to 512 protein nanopore channels charged by a voltage difference to generate electric currents. When a DNA strand passes through a nanopore, the nucleotides disrupt the electric current. The event type of the nucleotides passing through a nanopore can be determined by measuring the current flow. Approximately five nucleotides can remain in a single nanopore simultaneously; this affects the current signal with disruptions manifesting 4^5^ = 1024 different characteristics of all possible compound states.

To translate the characteristic disruptions into a DNA sequence, a caller is required to decode the bases (a.k.a *basecalling*). However, the fluctuations of the electric current are subtle and heavily influenced by noise due to the low signal-to-noise ratio (SNR) of the system. Furthermore, although the raw current signal is sampled at a constant time interval, the speed of a DNA strand passing through a nanopore is non-uniform. This leads to a change in the recording time of a single nucleotide. Hence, although MinION can rapidly generate long sequences, it suffers from low accuracy. While the read accuracy can easily surpass 99% for short reads with the current NGS technology, long reads from the nanopore sequencer still have an error rate of approximately 5% to 15%.

One of the solutions is duplex basecalling. By employing the template and complement strands simultaneously, duplex basecalling can provide significant improvement [1]. However, the data preparation requires special adapters and is not a typical attribute in most datasets. An alternative downstream method is to assemble multiple basecalled reads with overlaps to form a consensus sequence. This method requires large depth coverage, which cannot be achieved in certain applications [2]. Therefore, a basecalling algorithm providing a high read accuracy is essential in making the nanopore sequencer practical. There is an urgent demand for highly sophisticated basecallers.

In recent decades, deep neural networks (DNNs) have gained considerable attention in many research fields [3, 4, 5, 6, 7]. As DNNs are suitable for processing complicated problems involving big data, deep learning techniques are very promising for basecalling. Following this idea, official basecalling tools were developed by ONT, which include Metrichor, Albacore, Guppy, Scrappie, and Flappie, and Bonito. Most of these tools utilize DNN to achieve better results. However, Metrichor can only be executed online with cloud computing and was discontinued after 2017 [8]. Albacore is an earlier basecalling tool that can only run on CPUs. Guppy is designed as an improved version of Albacore and can be executed on GPUs. Scrappie is a *technology demonstrator* that aims to test the performance and robustness of newly developed software [9]. Currently, Scrappie has been replaced by Flappie. Flappie carries a connectionist temporal classification (CTC) [10] module as a decoder, leading to a segmentation-free method for base prediction. Recently, Bonito [11] provides another open-source python-based environment with outstanding performance. To evaluate various basecalling tools, Wick *et al*. [2] prepared a benchmark dataset by collecting bacterial genomes from the Proteobacteria phylum, with an emphasis on the Enterobacteriaceae family and Klebsiella pneumoniae.

In addition to official ONT basecalling tools, some researchers have released several third-party open-source basecallers. Similar to the function of Metrichor, Nanocall [12] is an offline open-source basecaller. It utilizes hidden Markov models [13] (HMMs) and models the events of all possible nucleotide combinations as hidden states. DeepNano [14] and DeepNano-blitz [15] employ bidirectional gated recurrent units (GRUs), a type of recurrent layer with fewer parameters [16]. SACall [17] combines the transformer self-attention layers [18] into a neural network, and outperforms most ONT official basecallers in eight out of nine bacterial genomes. It uses a CTC module as a decoder on top of their architecture, enabling the direct translation of raw current signals into DNA sequences.

As mentioned above, a nucleotide can be generated from a varying number of electrical signal measurements. In addition, segmentation of the characteristic disruption of the current signal can be difficult and harmful in identifying homopolymer segments. It may also cause a non-consecutive data processing flow and inevitably make the entire process incompatible with the neural network in terms of end-to-end training. Therefore, many studies have focused on aggregating dependent data to form a more performant feature.

In this paper, a novel basecalling neural network architecture called MSRCall is presented (Figure 1). The proposed MSRCall first uses convolutional layers to manipulate multi-scale downsampling. These back-to-back convolutional layers aim to capture features with receptive fields at different levels of complexity. As the moving speed of the DNA strand is inconsistent, it is difficult to select features of different ranges using fixed convolutional layers. A bidirectional LSTM [19, 20] was also adopted for recurrent layers to automatically estimate the importance of dependence in any range. Furthermore, the features of the RNN were aggregated at multiple scales with a fusion layer, followed by a CTC decoder at the rear end of the network to generate the final nucleotide outputs. To evaluate the performance of MSRCall, experiments were conducted using the dataset generated by [2]. The results obtained were the best from the perspective of read accuracy as well as consensus accuracy compared to Albacore, Guppy, DeepNano-blitz, and SACall.

**Figure 1:**
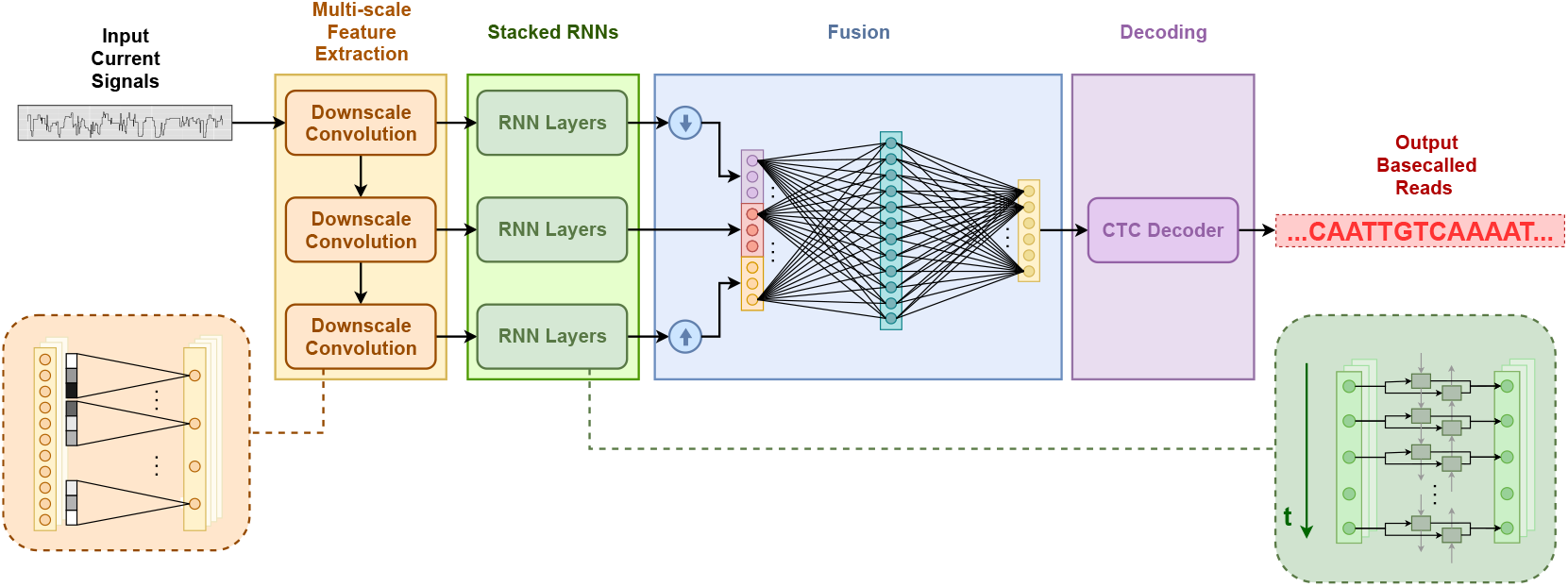
Overall architecture of the proposed MSRCall. It takes the squiggle current signals as input and outputs the decoded nucleotide predictions. The large orange block is the multi-scale structure; the large green block is the recurrent part; and the large blue block is the fusion block and the CTC decoder.

The main contribution of this work is a basecaller architecture design called MSRCall. It simultaneously utilizes multi-scale convolutional and bidirectional LSTM layers to capture semantic information. MSRCall is end-to-end trainable and does not require raw signal segmentation but adheres to the idea of segmenting signals at various levels, leading to more robust basecalling features. With multi-scale downsampling, bidirectional LSTM layers can focus on modeling the local dependencies under different receptive fields. The proposed architecture is more scalable when compared to transformer-based models. Compared to the state-of-the-art method, SACall, the MSRCall model can improve the read and consensus accuracy from 91.117% to 92.215% and Q25.5 (99.719%) to Q33.2 (99.952%), respectively.

The rest of this paper is organized as follows. The preprocessing and labeling of the adopted dataset, as well as the proposed architecture, are described in Sec. 2. The qualitative evaluations for several state-of-the-art basecallers and the proposed MSRCall are presented in Sec. 3. Sec. 4, concludes the paper.

## 2 Materials and methods

This section introduces the benchmark dataset and the design of our proposed MSRCall (multi-scale recurrent caller) network. After an initial description of the adopted dataset, where the trimming, preprocessing, and enhancement of the benchmark dataset are explained and evaluated, detailed descriptions and an analysis of the layer selections in the MSRCall model are provided along with an overview of the proposed MSRCall architecture.

### 2.1 Dataset and preprocessing

The training set has a total of 226,166 samples of DNA from 50 different isolate genomes, including 30 belonging to *Klebsiella pneumoniae*, 10 of Enterobacteriaceae, and 10 of Proteobacteria genomes. Wick *et al*. collected and released a massive dataset using MinION R9.4 flowcells [2]. Independent of the training set, this dataset also provides a testing set across bacteria from *Klebsiella pneumoniae, Shigella sonnei, Serratia marcescens, Haemophilus haemolyticus, Acinetobacter pittii, Stenotrophomonas maltophilia*, and *Staphylococcus aureus*. In addition, Wick *el al*. provided reference contig sequences for the testing set. These highly reliable contigs are constructed by assembling reads from Illumina using the SKESA assembler. Because this dataset is comprehensive and large enough for qualitative evaluation, it is used to evaluate the performance of neural network basecallers.

As the training data are not labeled, one approach is to obtain labels by directly applying ONT official basecalling tools such as Guppy. However, it has error rates ranging from 5% to 15%. As an alternative, we adopted a labeling method used in SACall [17] to collect better quality nucleotide labels.

Before the preprocessed current values X are used as training data for deep neural networks, it is common to normalize the distribution by subtracting the median value 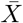 from the current value and dividing the result by the median absolute deviation σ_*X*_. That is:

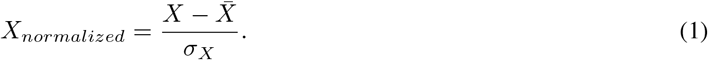

In general, the data normalization step can efficiently facilitate the convergence of deep neural networks. Finally, the normalized current signals and previously generated labels were used for network training.

### 2.2 Proposed architecture

As depicted earlier, the measurement process for MinION is quite complicated. ONT has released various kinds of pores with differing nanopore chemistries. Among all nanopore versions, R9.4 is currently the longest running product. Each measured current signal from the ONT sequencer implies that approximately five consecutive bases reside in the nanopore. The sequencer takes measurements at a rate of 4 kHz, and the DNA sequence moves at a velocity of 450 bases/s, producing an average of nine measurements for each base [21]. Although the DNA sequence is expected to pass through steadily, the DNA fragments may move at an inconsistent speed, causing the detected signal to either remain unchanged or skip more than one nucleotide. Besides, the nanopore current measurement is inevitably noisy and stochastic. Therefore, modeling the basecalling process is extraordinarily challenging. Because DNNs are effective at extracting robust features for complicated tasks, a novel deep structure called MSRCall was developed to disentangle the relationship between raw signal data and nucleotide labels.

The overall architecture is shown in Figure 1. Initially, there is a series of convolutional layers in the MSRCall model. These convolutional layers simultaneously extract low-level to high-level features, downsample the input length, and generate multi-scale receptive field data in the time domain. Because recurrent layers are capable of finding both local and global dependency, they are introduced to the network architecture to retrieve features missed by convolutional layers. In the MSRCall model, the recurrent layers were placed after the convolutional layers to reduce the computational complexity of the model. The resulting recurrent features are then adjusted to a uniform time scale. With two fully connected layers and one softmax layer appended in the following stage, it becomes possible to fuse and generate the probabilities of different events. Considering that the length of the calculated probabilities and the training labels are not always the same, the CTC [10] model is used to resolve the discrepancy.

Inspired by the research in natural language processing (NLP) [10], different basecalling studies have shown that it is beneficial to not force the exact mapping between the input signals and output labels [17]. Using a blank token, the CTC model can prevent basecalling redundant nucleotides from the same event while preserving the homopolymer cases. The output of the CTC module is the final predicted sequence of our MSRCall. The details of our network design are elaborated in the following subsections.

#### 2.2.1 Multi-scale convolutional layers

At the front end of the MSRCall architecture is the multi-scale structure. In practice, three convolutional layers [5] are stacked to generate multi-scale features. These convolutional layers are continuously connected sequentially, leading to patterns from low to high levels. Each convolutional layer downsamples signals in the time axis and simultaneously extends the channel depth of the features. To facilitate the discussion, it is initially assumed that an input signal 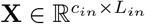 passing through a one-dimensional convolutional layer, where c_*in*_ is the channel number and L_*in*_ is the length of the input signal.

The forward calculation of the convolutional layer with c_*out*_ filters (and biases 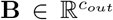) is actually the crosscorrelation of the input signal and every filter w_*i,j*_ ∈ ℝ^*k*^ with a window size k in weights 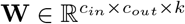. Formally, each channel of output features **Y** can be calculated as:

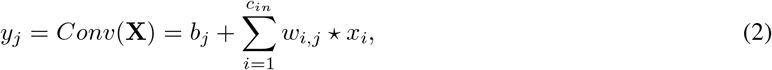

where c_*in*_ (c_*out*_) is the number of input (output) channels, and the symbol ⋆ denotes the cross-correlation operation. The length of y_*j*_ is:

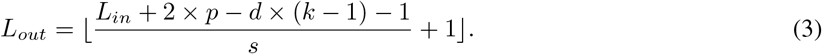

Here, p, d, k, and s are the sizes of padding, dilation, kernel window, and stride in the convolutional layer, respectively. The parameters p = 3, d = 1, k = 7, and s = 2 are set in all convolutional layers in the first stage of the model. As for the output channel size of each convolutional layer, the first layer outputs features of size c_*model*_/4, while the second and third produce features with c_*model*_/2 channels; c_*model*_ is the feature size of the entire model, which was set to 256 in our experiments.

In addition, a batch normalization [22] layer together with a rectified linear unit [23] (ReLU) layer were placed immediately after a convolutional layer. A batch normalization layer normalizes the input by subtracting the mean of the mini-batch and then dividing the input by the standard deviation of the mini-batch. The precise calculation of a batch normalization layer is as follows:

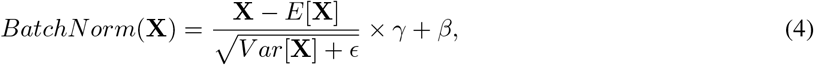

where ϵ, γ, and *β* are learnable parameters of the batch normalization layer.

A batch normalization layer strives to maintain the input at a zero mean and a standard deviation of one. In practice, batch normalization helps avoid internal covariate shift, which is a severe problem that can cause limited learning efficiency, overfitting, and gradient vanishing (or exploding).

The ReLU layer is a conventional activation function that is widely used after a convolutional layer. It is a non-linear transform that trims negative values to zero. A non-linear operation helps separate convolutional layers and consequently increases model feasibility.

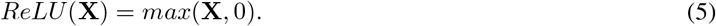

### 2.2.2 Recurrent layers

To best encode the sequence data, recurrent layers are attached after the convolutional features for each scale. Popular recurrent layers include LSTM [20], GRU [16], and the transformer layer [18]. These layers are widely used and proven to be stable in a variety of applications in NLP, which is also a task for analyzing time-domain signals. Unlike the ordinary feedforward layer, an LSTM layer has a time-crossing feedback connection, which is helpful in capturing relations in a time series. Benefiting from the memory structure, LSTM layers are capable of learning long-term dependency with resistance to the vanishing gradient problem encountered by vanilla RNN layers. GRU is a simplified version of an LSTM layer, and has a faster inference speed than an LSTM layer, with a trade-off in performance. The design of the transformer layer is entirely different from that of previous recurrent structures. Based on an attention mechanism, a transformer layer is more suitable for parallel computing. Empirically, transformer layers are faster in very long queries, whereas recurrent layers are faster in shorter queries.

By contrast, parameter tuning in transformer layers is more challenging than in recurrent layers and mostly requires pretraining to achieve the best outcome, which is impractical in the basecalling problem. Furthermore, when basecalling nanopore sequencing data, adjacent signals are much more relevant than those separated by an appreciable distance.

As a result, an improved version of LSTM, the bidirectional LSTM layer [19], is selected to extract sequencing features. The design of the LSTM layer, which naturally tends to discard information that is spread out, is more suitable for basecalling. There are five consecutive bidirectional LSTM layers in our MSRCall model. The bidirectional LSTM structure consists of two LSTM layers: one layer takes input sequence data along the positive time direction and the other takes data along the negative time direction [19]. Through this mechanism, it becomes possible to keep the information on forward and backward states together. The channel size for each LSTM feature was set to 128 in our model.

The following are the forward direction behavior summaries of the bidirectional LSTM layer.

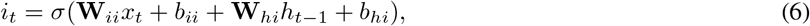

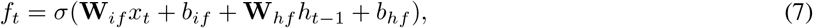

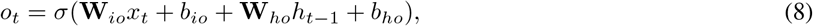

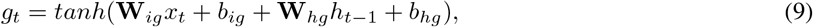

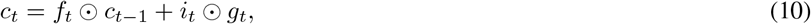

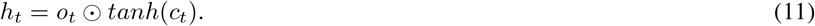

Equations (6) to (9) are the calculations for the four gates in an LSTM layer: input gate, forget gate, output gate, and modulation gate, where different *b* and **W** are the biases and weights, respectively, for the input x_*t*_ and hidden states *h*_*t*_ −_1_ of the four gates; *c*_*t*_ in Equation (10) denotes the memory cell. The Hadamard operation⊙ is used in c_*t*_ to calculate the output product of the modulation gate (*g*_*t*_) and the input gate (*i*_*t*_), as well as the product of the hidden state at time *t* −1 (*h*_*t*_− _1_) and the output of the forget gate (*f*_*g*_). Finally, *c*_*t*_ is controlled by the output gate o_*t*_, and thus generates the final result in the forward direction, *h*_*t*_, at time step t. The behaviors for the backward direction in the bidirectional LSTM layer can be obtained by reversing the input signals in the above equations.

Instead of applying the batch normalization method introduced in Sec. 2.2.1, the method suggested in this work is to couple the LSTM layers with layer normalization [24]. The key concept of layer normalization is borrowed from batch normalization. While batch normalization computes across the mini-batch, layer normalization normalizes the input across the feature depth dimensions. Empirically, it is more robust and stable to adopt layer normalization after a recurrent layer [25].

Following the LSTM layers, there are re-scaling layers and a fusion layer to merge recurrent features from every scale. A feature-pooling layer (a 1× 1 convolutional layer in our design) is adopted for the fusion function. After all re-scaled features are concatenated, the feature pooling layer produces an integrated feature with a channel depth of c_*model*_ = 256.

#### 2.2.3 CTC decoder

Once the robust multi-level features for base prediction are obtained, the final step is to adjust the feature size to fit the ground truth labels and apply a loss function to make the entire neural network trainable. Then, the results in the previous stage are blended with a fully connected layer and a softmax layer. These two layers collectively act as a classifier to predict probabilities for nucleotides. The combination of the fully connected layer and the softmax layer performs the following calculation:

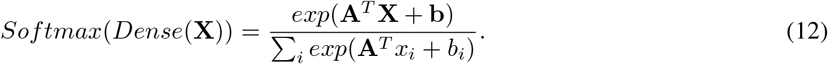

The *Dense*(.) operation indicates a fully connected layer that is equivalent to a linear transform with weights W and biases b. The softmax layer normalizes the output vector. All components in the vector are mapped into the interval (0, 1), with their sum equal to 1. Therefore, the result of the softmax layer can represent the probability distribution over the predicted output classes.

The prediction is decoded using the CTC model, which has been demonstrated to be superior in various sequence-to-sequence [26] tasks such as speech recognition [27] and machine translation [28]. Although input and output alignment is not required for the CTC algorithm, the mapping is restricted to the many-to-one scenario. The most prominent feature of the CTC model is steady management when the outcome has two (or more) identical characters in a row. These properties are in exact agreement with basecalling nanopore current signals, especially for homopolymer repeats.

The space of the output symbols can be set to {*A, T, C, G, ϵ, ϕ*}.The CTC module treats the ϵ token as a separator for two identical valid characters. Other repeating characters without the ϵ token will collapse during the decoding process. In other words, the decoded sequences of *AATTC* and *AϵATϵTC* would be *ATC* and *AATTC*, respectively.

During training, a CTC loss function [10] accumulates the probability of output predictions and then calculates the edit distance to the reference DNA sequence. As for the testing phase, all possible output sequences can be exhaustively enumerated. However, this is impractical when the sequence is very long, and an enormous number of combinations exist. Hence, the beam search algorithm is often adopted for the inference stage. The beam search algorithm is a trade-off method between greedy and exhaustive searches. In the beam search process, the paths taken at a time step are those with the highest k scores, while k (set to 3 in our experiments) is the beam width. Based on these results, the paths with the top k scores are determined at the next time step. This process is iterated until the last time step.

## 3 Results

This section presents, from various perspectives, the standard metrics that are widely used when assessing a basecaller. In cases of data having a low read depth, as in clinical metagenomics, there may be insufficient repeated information to correct the mistaken loci. Therefore, applications with a low data depth generally emphasize read identity (read accuracy). In contrast, for applications with a high read depth, high-quality sequences can be constructed through the assembling process. In these applications, the focus is more on the correctness of the final assemblies rather than the precision of every individual read, making consensus accuracy the preferred metric.

Experiments were conducted on the dataset collected by Wick *et al*. [2] Initially, the accuracy and error type on read level was investigated. Next, the quality of the assembly level was examined from various perspectives, including statistics such as N50 and consensus accuracy. The model was then compared with Bonito’s [11] on the open-source dataset. The proposed MSRCall model was also examined on another testing set that includes *Lambda phage, E. coli*, human, and *Klebsiella pneumoniae*. Finally, an ablation study was carried out to verify the effectiveness of multi-scale structure. The bidirectional LSTM was selected for the recurrent layer instead of the transformer layer because the former has better performance. (Supplementary Table S1)

### 3.1 Read identity

In this paper, the read accuracy refers to the BLAST identity [30]. First, the basecalled result of each read is aligned to the reference genome using minimap2 [31]. The accuracy of each read is the number of matching bases divided by the total alignment length, including insertion, deletion, match, and mismatch bases. Following standard procedure, 0% accuracy was assigned to the defective basecalled reads using 50% accuracy as the threshold. Then, the median value was used to determine the overall read accuracy of all reads, including those of defective reads. The overall read accuracy comparison of each method is presented in Table 1. The results show that the proposed MSRCall achieved state-of-the-art read accuracy for every bacterial genome being tested. In terms of read accuracy, MSRCall outperformed SACall by 1.1% on average.

**Table 1:**
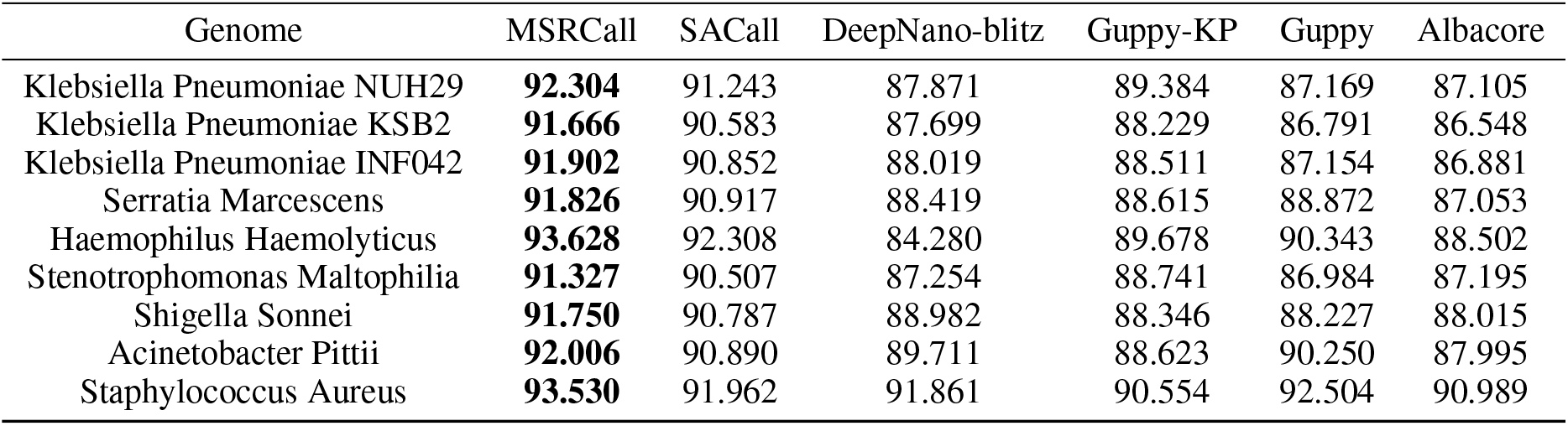
Read accuracy (%) of testing different bacterial genomes from the dataset [2]. The highest read accuracy is emphasized in bold for each isolated genome.

| Genome | MSRCall | SACall | DeepNano-blitz | Guppy-KP | Guppy | Albacore |
| --- | --- | --- | --- | --- | --- | --- |
| Klebsiella Pneumoniae NUH29 | **92.304** | 91.243 | 87.871 | 89.384 | 87.169 | 87.105 |
| Klebsiella Pneumoniae KSB2 | **91.666** | 90.583 | 87.699 | 88.229 | 86.791 | 86.548 |
| Klebsiella Pneumoniae INF042 | **91.902** | 90.852 | 88.019 | 88.511 | 87.154 | 86.881 |
| Serratia Marcescens | **91.826** | 90.917 | 88.419 | 88.615 | 88.872 | 87.053 |
| Haemophilus Haemolyticus | **93.628** | 92.308 | 84.280 | 89.678 | 90.343 | 88.502 |
| Stenotrophomonas Maltophilia | **91.327** | 90.507 | 87.254 | 88.741 | 86.984 | 87.195 |
| Shigella Sonnei | **91.750** | 90.787 | 88.982 | 88.346 | 88.227 | 88.015 |
| Acinetobacter Pittii | **92.006** | 90.890 | 89.711 | 88.623 | 90.250 | 87.995 |
| Staphylococcus Aureus | **93.530** | 91.962 | 91.861 | 90.554 | 92.504 | 90.989 |

In addition, a comprehensive analysis on the errors of the basecalled reads is reported in the Supplementary Table S3. The MSRCall deletion, insertion, mismatch, and total error rates are the lowest among the different methods. The only exception is the deletion rate for *Haemophilus haemolyticus*, which is merely 0.05% higher than that of Albacore.

### 3.2 Assembly error profiles

It should be noted that a high read identity does not always imply good assembly results. Although random errors can be removed statistically with repeated information covered by high read depth, systematic errors at the same locus can still appear across the reads. Therefore, even with a high read depth and high read accuracy, the assembly can still be imperfect because of systematic errors. Following the evaluation protocol of [17], consensus sequences were generated by gathering the basecalled read sets. Then, the resulting assembly contigs were assessed by Quast [29], a tool for evaluating genome assemblies. The consensus identity, number of misassemblies, mismatches, indels, N50, and genome fractions are reported in Table 2 (For the comprehensive table, see Supplementary Table S4).

**Table 2:**
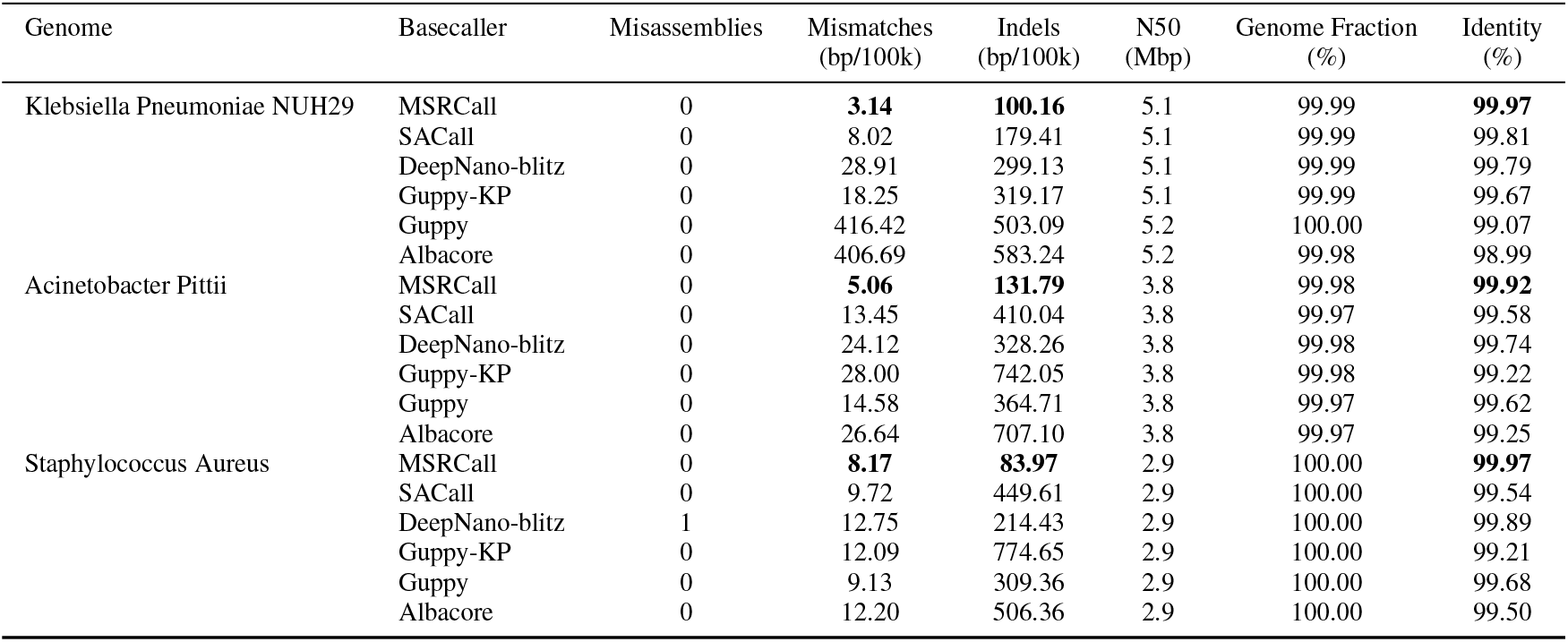
Consensus analysis of all nine bacterial genomes being tested from the benchmark dataset [2] for five different basecalling models. In addition to consensus identity, analytic reports on all assemblies were obtained using Quast [29].

The proposed MSRCall outperforms all novel methods in almost every perspective in the Quast summary. Although one more misassembly was obtained compared with SACall, Guppy, and Albacore on the genome sequence reads from the isolate of *Serratia Marcescens*, the best consensus accuracy was still obtained for *Serratia Marcescens*. The number of normalized base pairs counting mismatches and *Indel* (insertion plus deletion) were significantly reduced compared to all other methods, resulting in a considerably higher accuracy for all isolate genomes tested.

Figure 2 provides the Phred quality score (Qscore) representation for all assemblies. The definition of Qscore is given by

**Figure 2:**
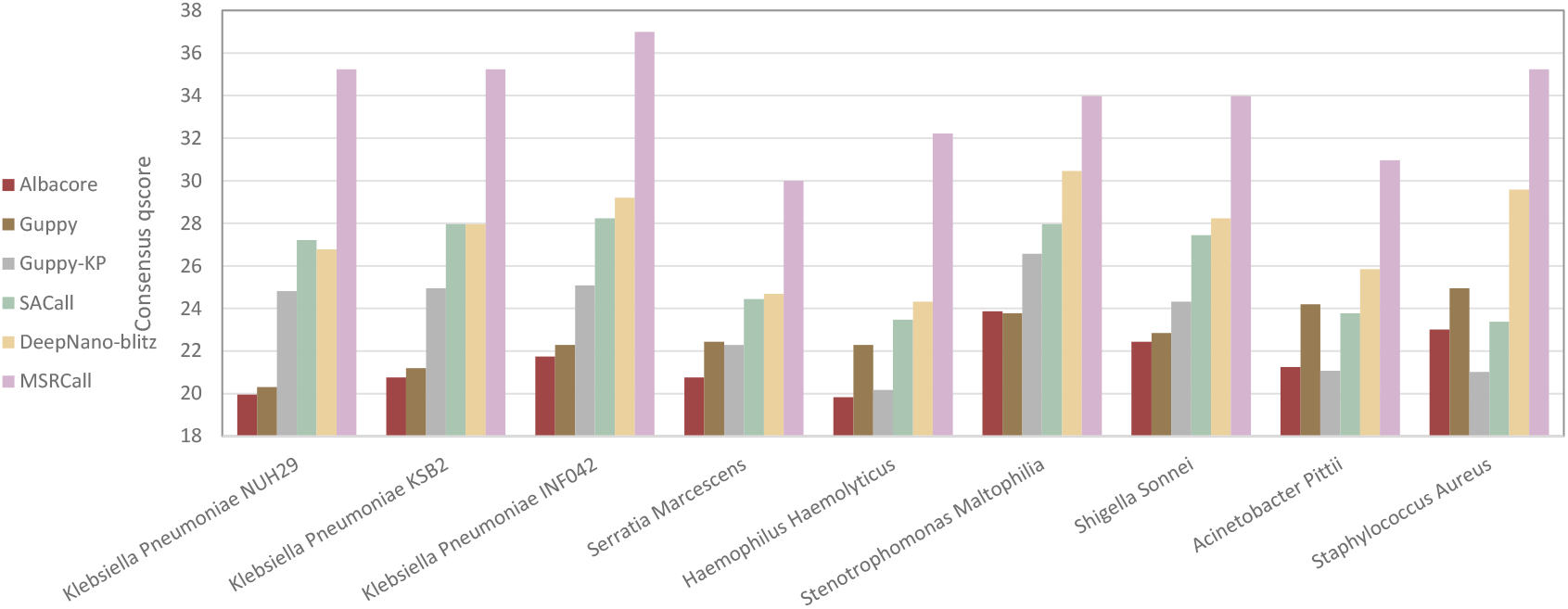
Phred quality score (qscore, higher is better) plot of consensus identity for five different basecallers on the testing set.

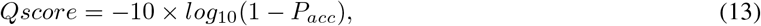

where P_*acc*_ is the average accuracy in the range [0, 1]. From the viewpoint of the mean consensus Qscore, our MSRCall model improves from +7.70 to +11.88.

### 3.3 Comparison with more methods

Recently, the latest Oxford Nanopore Technologies’ basecaller, Bonito [11], achieved outstanding performance. However, the dataset used for Bonito’s pretrained model is not publically available. In addition to data preprocessing and postprocessing, the dataset released by Bonito was used to retrain Bonito and the MSRcall model of this study, for a fair comparison. The two models were also evaluated on the testing set from Wick’s [2] with the output length set to 4096. In Table 3, it is observed that the proposed method outperforms Bonito’s model. The detailed analysis of consensus errors was also examined. MSRCall outperformed for most of the bacterial genomes shown in Figure 3. In addition, the MSRCall model was tested on the testing set used in [32] to verify the robustness of the model across different datasets. This testing set includes *Lambda phage, E. coli*, human, and *Klebsiella pneumoniae*. The MSRCall model achieved, on average, a 3.9% higher read identity than CausalCall on the testing sets (reported in Supplementary Table S2).

**Table 3:**
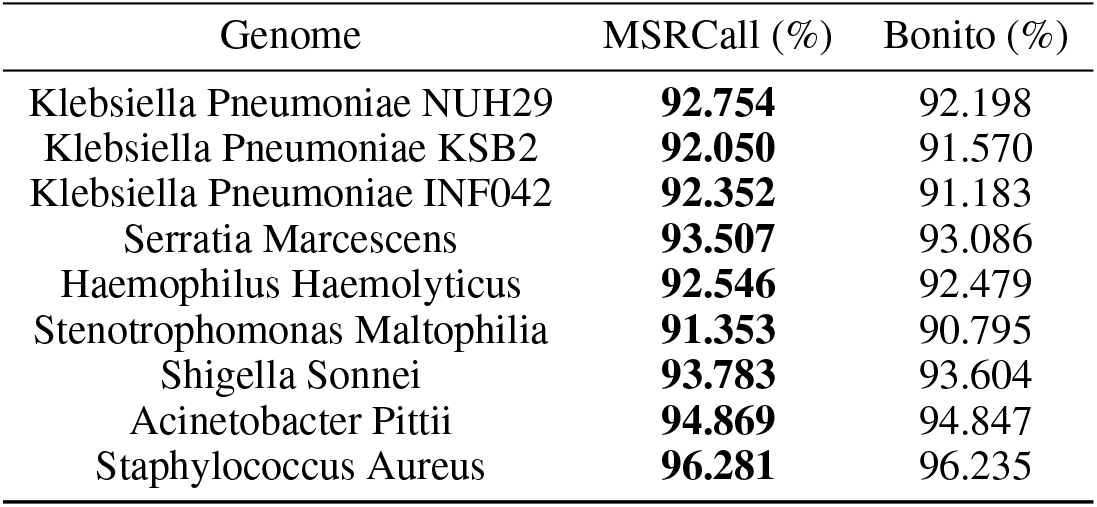
Read accuracy for MSRCall and Bonito models trained on the dataset provided by Bonito.

| Genome | MSRCall (%) | Bonito (%) |
| --- | --- | --- |
| Klebsiella Pneumoniae NUH29 | **92.754** | 92.198 |
| Klebsiella Pneumoniae KSB2 | **92.050** | 91.570 |
| Klebsiella Pneumoniae INF042 | **92.352** | 91.183 |
| Serratia Marcescens | **93.507** | 93.086 |
| Haemophilus Haemolyticus | **92.546** | 92.479 |
| Stenotrophomonas Maltophilia | **91.353** | 90.795 |
| Shigella Sonnei | **93.783** | 93.604 |
| Acinetobacter Pittii | **94.869** | 94.847 |
| Staphylococcus Aureus | **96.281** | 96.235 |

**Figure 3:**
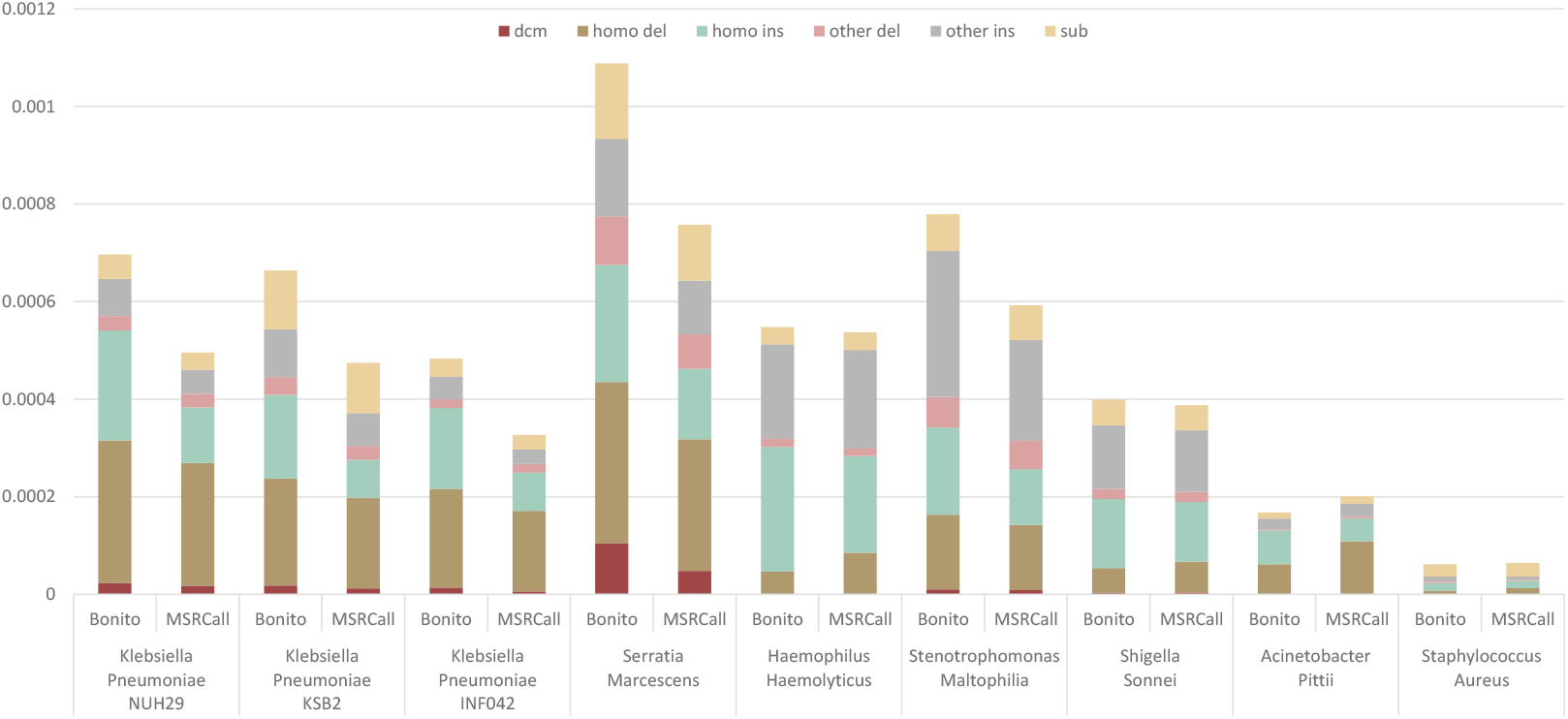
Detailed analysis of consensus errors for Bonito and our proposed model. Error types include Dcmmethylation (dcm), homopolymer insertion (homo ins), homopolymer deletion (homo del), other insertions (other ins), other deletions (other del), and substitutions (sub).

### 3.4 Ablation study

To better disentangle the actual effect of multi-scale structure, an ablation experiment was conducted. The performance of the MSRCall model was assessed with different numbers of scales. As observed in Table 4, the performance degrades, as expected, as the number of scales decreases.

**Table 4:** Read accuracy comparison for MSRCall models with various number of scales.

| Genome | 3 scales (%) | 2 scales (%) | 1 scale (%) |
| --- | --- | --- | --- |
| Klebsiella Pneumoniae NUH29 | **92.304** | 90.635 | 90.316 |
| Klebsiella Pneumoniae KSB2 | **91.666** | 89.923 | 89.674 |
| Klebsiella Pneumoniae INF042 | **91.902** | 90.173 | 89.924 |
| Serratia Marcescens | **91.826** | 90.424 | 89.933 |
| Haemophilus Haemolyticus | **93.628** | 91.618 | 91.314 |
| Stenotrophomonas Maltophilia | **91.327** | 89.852 | 89.537 |
| Shigella Sonnei | **91.750** | 90.057 | 89.921 |
| Acinetobacter Pittii | **92.006** | 90.343 | 89.897 |
| Staphylococcus Aureus | **93.530** | 91.759 | 91.228 |

## 4. Conclusion

In this work, a novel multi-scale recurrent basecaller architecture, namely MSRCall, was proposed. To utilize the nature of the nanopore signals, recurrent features of different receptive fields were collected. Three-stage convolutional layers were adopted to perform downsampling on the time axis before applying the bidirectional LSTM layers. The resulting multi-scale temporal features were more robust to the speed fluctuation of nucleotides, which modulate the electrical signals collected by ONT’s MinION sequencers. The recurrent features were further fused and the CTC loss was utilized to train the entire network. Finally, the MSRCall model was trained on a publicly available dataset and the results were compared with those of other state-of-the-art methods on a testing set. The proposed MSRCall provided distinctive improvements on both read accuracy and consensus accuracy. Compared to the popular SACall basecaller, the MSRCall was able to enhance the average read accuracy from 91.117% to 92.215% and consensus accuracy from 99.719% to 99.952%.

