## Supplementary materials for "MSRCall: A Multi-scale Deep Neural Network to Basecall Oxford Nanopore Sequences"

### 1 Selection of recurrent neural networks in our model

The transformer layer is known for its high parallelizability in the encoding stage. Consequently, it is sometimes considered a better choice for a recurrent layer. However, it also suffers from the following practical problems: first, it is difficult to have a transformer layer converge without applying pretrained weights, which is not available for basecalling; further, the positional encoding mechanism is relatively weak in maintaining the correct order, which is critical in DNA sequencing.

Although transformer layers have been used in SACall [1], LSTM was adopted in this work. To demonstrate that LSTM is the right choice for the recurrent layer in the MSRCall model, a second MSRCall architecture was constructed to replace LSTM with transformer layers. This model was designed with layer parameters close to the LSTM-based version and was slightly larger in model size (approximately $27\%$ larger). The difference in read accuracy is presented in Table S1. The results show that using transformer layers is less effective in this situation. Moreover, even with the maximum available batch size on the same device, the NVIDIA RTX 2080 Ti GPU, the transformer-based model can only run at a speed that is $34.3\%$ slower than the LSTM-based version.

### 2 Training scheme

Our experiments were conducted on the dataset collected by Wick *et al.* [2]. To objectively evaluate the proposed MSRCall, our results were compared with those of Albacore, Guppy, Guppy-KP, and SACall.

In the training phase, the Adam optimizer [3, 4] was applied to backpropagate the loss and update the parameters of the MSRCall. The learning rate was set to $0.001$, and the weight decay was $0.01$. The coefficients $\beta_1$ and $\beta_2$, used to calculate the running averages of the gradient and its square, were set to $0.9$ and $0.999$, respectively.

### 3 Consensus error analysis

To better understand the various types of consensus basecalling errors, the post-processing flow suggested by the official ONT group was followed to polish the assemblies in the previous stage with four rounds of Racon [5] and one round of Medaka [6]. Racon is a polishing tool used after assembly to improve the quality of raw uncorrected contigs. Medaka is another tool that can create consensus sequences from nanopore sequencing data using neural networks. The results were compared with those of SACall in Figure S1.

Table S1: Read accuracy comparison for MSRCall models with transformer and bidirectional LSTM layers.

| Genome | Bi-LSTM (%) | Transformer (%) |
|---|---|---|
| Klebsiella Pneumoniae NUH29 | **92.304** | 87.285 |
| Klebsiella Pneumoniae KSB2 | **91.666** | 86.613 |
| Klebsiella Pneumoniae INF042 | **91.902** | 86.957 |
| Serratia Marcescens | **91.826** | 87.251 |
| Haemophilus Haemolyticus | **93.628** | 86.953 |
| Stenotrophomonas Maltophilia | **91.327** | 86.962 |
| Shigella Sonnei | **91.750** | 86.707 |
| Acinetobacter Pittii | **92.006** | 86.654 |
| Staphylococcus Aureus | **93.530** | 86.646 |

Table S2: Read accuracy for MSRCall and CausalCall models testing on the datasets used in [7].

| Genome | MSRCall (%) | CausalCall (%) |
|---|---|---|
| Lambda phage | **93.075** | 89.21 |
| E. coli | **94.616** | 89.48 |
| Human | **89.243** | 86.88 |
| K. pneumoniae | **92.358** | 88.12 |

The consensus errors were evaluated by breaking down the errors into six different error sources: Dcm-methylations, homopolymer insertions, homopolymer deletions, other insertions, other deletions, and substitutions. Dcm-methylations are the errors that occur in the CCAGG/CCTGG Dcm motif. Homopolymer errors, including homopolymer insertions and deletions, refer to incorrect length predictions in homopolymer repeat segments with three or more bases in the reference sequence. The insertion and deletion errors that occur at non-homopolymer loci are considered to be other insertions and deletions. The substitution errors are the mismatch bases between the contigs and the reference sequence. Medaka (version 1.2.1) was adopted in our flow and SACall was reran with this newer version for a fair comparison.

The diagram in Figure S1 shows that the incorrect homopolymer length is the most dominant error type, which takes up 78.65% and 68.30% of total errors on average for SACall and MSRCall, respectively. It is also observed that almost all variants of the errors are significantly reduced. The only error type that slightly increased in our model was the homopolymer insertion. However, when the overall incorrect homopolymer length (i.e., both homopolymer insertion plus deletion) is considered, the homopolymer errors are reduced significantly, ranging from 51.4% in *Stenotrophomonas Maltophilia* to 80.8% in *Staphylococcus Aureus*, compared to SACall. The results indicate that homopolymer calls are fundamentally improved with the proposed basecaller.

[4] Ilya Loshchilov and Frank Hutter. Decoupled weight decay regularization. *arXiv preprint arXiv:1711.05101*, 2017.

[5] Isovic. isovic/racon, 2017.

[6] Nanoporetech. nanoporetech/medaka, 2017.

[7] Jingwen Zeng, Hongmin Cai, Hong Peng, Haiyan Wang, Yue Zhang, and Tatsuya Akutsu. Causalcall: Nanopore basecalling using a temporal convolutional network. *Frontiers in Genetics*, 10:1332, 2020.

[8] Alexey Gurevich, Vladislav Saveliev, Nikolay Vyahhi, and Glenn Tesler. Quast: quality assessment tool for genome assemblies. *Bioinformatics*, 29(8):1072–1075, 2013.


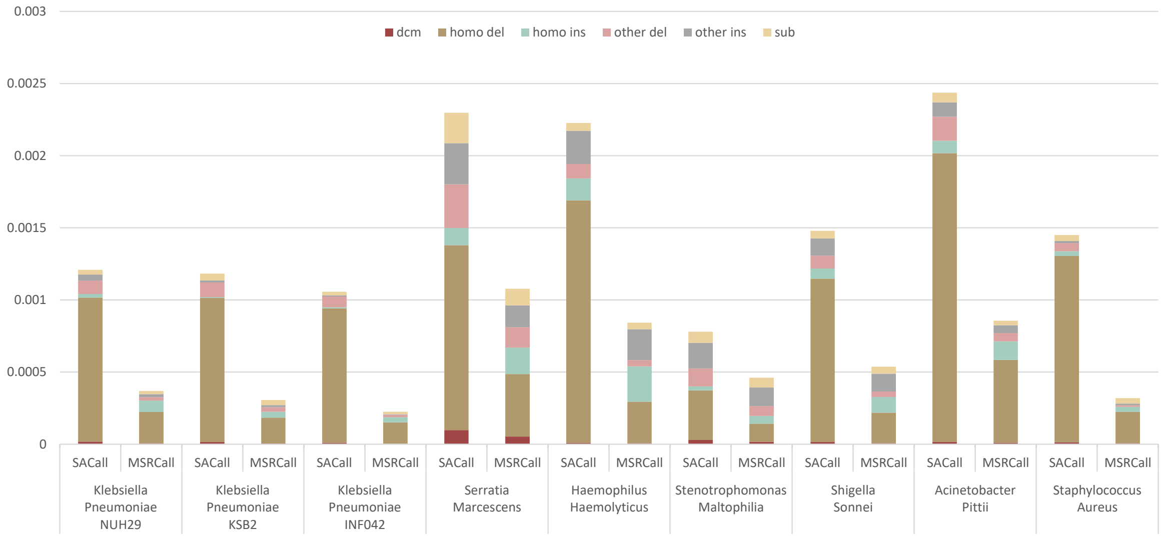


Figure S1: Detailed analysis of consensus errors for SACall and our proposed model. Error types include Dcm-methylation, homopolymer insertion, homopolymer deletion, other insertions, other deletions, and substitutions.

Table S3: Error analysis of tests performed on different bacterial genomes from the benchmark dataset [2]. The bold text indicates the lowest error rate for each category across the five basecallers. Here, the definitions of deletion rate, insertion rate, and mismatch rate are the number of deletions, insertions, and mismatches over the total alignment length, respectively.

| Genome | Basecaller | Deletion rate(%) | Insertion rate(%) | Mismatch rate(%) | Error rate(%) |
|---|---|---|---|---|---|
| Klebsiella Pneumoniae NUH29 | MSRCall | **4.45** | **2.40** | **3.07** | **9.91** |
| | SACall | 4.93 | 2.63 | 3.50 | 11.06 |
| | DeepNano-blitz | 5.88 | 3.70 | 4.69 | 14.26 |
| | Guppy-KP | 5.68 | 3.33 | 4.09 | 13.10 |
| | Guppy | 4.85 | 5.51 | 5.34 | 15.69 |
| | Albacore | 4.85 | 5.76 | 5.08 | 15.69 |
| Klebsiella Pneumoniae KSB2 | MSRCall | **4.58** | **2.21** | **3.30** | **10.10** |
| | SACall | 5.26 | 2.31 | 3.69 | 11.26 |
| | DeepNano-blitz | 6.00 | 3.25 | 4.87 | 14.13 |
| | Guppy-KP | 5.90 | 3.38 | 4.56 | 13.84 |
| | Guppy | 5.42 | 4.65 | 5.66 | 15.73 |
| | Albacore | 5.37 | 4.99 | 5.45 | 15.80 |
| Klebsiella Pneumoniae INF042 | MSRCall | **4.39** | **2.17** | **3.18** | **9.75** |
| | SACall | 5.13 | 2.27 | 3.56 | 10.95 |
| | DeepNano-blitz | 5.82 | 3.24 | 4.75 | 13.81 |
| | Guppy-KP | 5.67 | 3.34 | 4.42 | 13.44 |
| | Guppy | 5.27 | 4.60 | 5.48 | 15.35 |
| | Albacore | 5.18 | 5.00 | 5.30 | 15.48 |
| Serratia Marcescens | MSRCall | **4.03** | **2.29** | **3.34** | **9.65** |
| | SACall | 4.37 | 2.42 | 3.72 | 10.51 |
| | DeepNano-blitz | 5.31 | 3.01 | 4.64 | 12.95 |
| | Guppy-KP | 5.21 | 3.28 | 4.40 | 12.90 |
| | Guppy | 4.47 | 3.56 | 4.56 | 12.59 |
| | Albacore | 4.50 | 4.87 | 5.18 | 14.55 |
| Haemophilus Haemolyticus | MSRCall | 3.28 | **2.44** | **2.70** | **8.43** |
| | SACall | 3.87 | 2.59 | 3.25 | 9.71 |
| | DeepNano-blitz | 5.70 | 3.46 | 4.89 | 14.05 |
| | Guppy-KP | 4.56 | 3.79 | 4.21 | 12.56 |
| | Guppy | 3.57 | 4.52 | 4.05 | 12.14 |
| | Albacore | **3.23** | 6.40 | 4.30 | 13.93 |
| Stenotrophomonas Maltophilia | MSRCall | **4.89** | **2.36** | **3.29** | **10.53** |
| | SACall | 5.20 | 2.60 | 3.62 | 11.42 |
| | DeepNano-blitz | 5.97 | 3.73 | 4.97 | 14.67 |
| | Guppy-KP | 5.90 | 3.28 | 4.12 | 13.30 |
| | Guppy | 5.08 | 5.10 | 5.18 | 15.36 |
| | Albacore | 4.98 | 5.40 | 4.85 | 15.24 |
| Shigella Sonnei | MSRCall | **4.48** | **2.06** | **3.57** | **10.11** |
| | SACall | 5.27 | 2.20 | 3.99 | 11.47 |
| | DeepNano-blitz | 5.66 | 2.47 | 4.65 | 12.78 |
| | Guppy-KP | 5.76 | 3.21 | 5.05 | 14.03 |
| | Guppy | 5.80 | 3.12 | 5.31 | 14.23 |
| | Albacore | 5.98 | 3.46 | 5.39 | 14.84 |
| Acinetobacter Pittii | MSRCall | **3.96** | **2.37** | **3.40** | **9.72** |
| | SACall | 4.87 | 2.43 | 3.94 | 11.23 |
| | DeepNano-blitz | 4.92 | 2.85 | 4.30 | 12.07 |
| | Guppy-KP | 5.59 | 3.38 | 4.61 | 13.58 |
| | Guppy | 4.45 | 3.38 | 4.35 | 12.18 |
| | Albacore | 4.52 | 4.91 | 5.01 | 14.44 |
| Staphylococcus Aureus | MSRCall | **3.41** | **1.84** | **2.44** | **7.69** |
| | SACall | 4.17 | 2.03 | 3.02 | 9.23 |
| | DeepNano-blitz | 4.22 | 2.08 | 3.12 | 9.42 |
| | Guppy-KP | 4.62 | 2.76 | 3.47 | 10.85 |
| | Guppy | 3.73 | 2.27 | 2.88 | 8.87 |
| | Albacore | 4.12 | 3.11 | 3.40 | 10.64 |

Table S4: Consensus analysis of all nine bacterial genomes being tested from the benchmark dataset [2] for five different basecalling models. Besides the consensus identity, analytic reports of all assemblies are performed using Quast [8]

| Genome | Basecaller | Misassemblies | Mismatches (bp/100k) | Indels (bp/100k) | N50 (Mbp) | Genome Fraction (%) | Identity (%) |
|---|---|---|---|---|---|---|---|
| Klebsiella Pneumoniae NUH29 | MSRCall | 0 | **3.14** | **100.16** | 5.1 | 99.99 | **99.97** |
| | SACall | 0 | 8.02 | 179.41 | 5.1 | 99.99 | 99.81 |
| | DeepNano-blitz | 0 | 28.91 | 299.13 | 5.1 | 99.99 | 99.79 |
| | Guppy-KP | 0 | 18.25 | 319.17 | 5.1 | 99.99 | 99.67 |
| | Guppy | 0 | 416.42 | 503.09 | 5.2 | 100.00 | 99.07 |
| | Albacore | 0 | 406.69 | 583.24 | 5.2 | 99.98 | 98.99 |
| Klebsiella Pneumoniae KSB2 | MSRCall | 0 | **4.88** | **111.23** | 5.2 | 99.99 | **99.97** |
| | SACall | 0 | 8.87 | 146.22 | 5.2 | 99.99 | 99.84 |
| | DeepNano-blitz | 0 | 33.20 | 321.29 | 5.2 | 99.99 | 99.84 |
| | Guppy-KP | 0 | 23.48 | 293.52 | 5.2 | 100.00 | 99.68 |
| | Guppy | 0 | 428.95 | 322.17 | 5.2 | 99.99 | 99.24 |
| | Albacore | 1 | 421.19 | 410.68 | 5.2 | 99.95 | 99.16 |
| Klebsiella Pneumoniae INF042 | MSRCall | 0 | **2.87** | **104.83** | 5.3 | 99.99 | **99.98** |
| | SACall | 0 | 5.68 | 146.46 | 5.3 | 100.00 | 99.85 |
| | DeepNano-blitz | 0 | 22.22 | 294.13 | 5.3 | 100.00 | 99.88 |
| | Guppy-KP | 0 | 18.06 | 292.23 | 5.4 | 100.00 | 99.69 |
| | Guppy | 0 | 255.27 | 327.19 | 5.3 | 100.00 | 99.41 |
| | Albacore | 0 | 255.79 | 410.21 | 5.4 | 100.00 | 99.33 |
| Serratia Marcescens | MSRCall | 2 | **19.41** | **125.48** | 5.5 | 99.24 | **99.90** |
| | SACall | 1 | 43.71 | 337.58 | 5.5 | 99.92 | 99.64 |
| | DeepNano-blitz | 1 | 88.03 | 339.19 | 5.5 | 99.96 | 99.66 |
| | Guppy-KP | 4 | 87.86 | 502.04 | 5.5 | 99.24 | 99.41 |
| | Guppy | 1 | 120.74 | 444.11 | 5.5 | 99.41 | 99.43 |
| | Albacore | 1 | 116.76 | 712.57 | 5.5 | 99.25 | 99.16 |
| Haemophilus Haemolyticus | MSRCall | 3 | **12.34** | **115.74** | 2.1 | 100.00 | **99.94** |
| | SACall | 3 | 14.88 | 428.25 | 2.1 | 100.00 | 99.55 |
| | DeepNano-blitz | 2 | 31.63 | 454.64 | 2.0 | 100.00 | 99.63 |
| | Guppy-KP | 3 | 24.48 | 917.01 | 2.1 | 100.00 | 99.04 |
| | Guppy | 3 | 16.40 | 556.48 | 2.1 | 100.00 | 99.41 |
| | Albacore | 3 | 22.91 | 956.52 | 2.1 | 100.00 | 98.96 |
| Stenotrophomonas Maltophilia | MSRCall | 0 | **11.26** | **74.31** | 4.8 | 100.00 | **99.96** |
| | SACall | 0 | 14.68 | 141.11 | 4.8 | 100.00 | 99.84 |
| | DeepNano-blitz | 0 | 25.15 | 155.66 | 4.8 | 100.00 | 99.91 |
| | Guppy-KP | 0 | 26.57 | 199.18 | 4.8 | 100.00 | 99.78 |
| | Guppy | 0 | 41.31 | 366.69 | 4.8 | 100.00 | 99.58 |
| | Albacore | 0 | 23.99 | 376.22 | 4.8 | 100.00 | 99.59 |
| Shigella Sonnei | MSRCall | 0 | **7.14** | **130.06** | 4.8 | 100.00 | **99.96** |
| | SACall | 0 | 9.21 | 171.07 | 4.8 | 100.00 | 99.82 |
| | DeepNano-blitz | 1 | 23.14 | 308.35 | 4.8 | 99.98 | 99.85 |
| | Guppy-KP | 0 | 21.66 | 351.57 | 4.8 | 100.00 | 99.63 |
| | Guppy | 0 | 290.03 | 227.08 | 4.8 | 100.00 | 99.48 |
| | Albacore | 0 | 287.69 | 274.00 | 4.8 | 99.99 | 99.43 |
| Acinetobacter Pittii | MSRCall | 0 | **5.06** | **131.79** | 3.8 | 99.98 | **99.92** |
| | SACall | 0 | 13.45 | 410.04 | 3.8 | 99.97 | 99.58 |
| | DeepNano-blitz | 0 | 24.12 | 328.26 | 3.8 | 99.98 | 99.74 |
| | Guppy-KP | 0 | 28.00 | 742.05 | 3.8 | 99.98 | 99.22 |
| | Guppy | 0 | 14.58 | 364.71 | 3.8 | 99.97 | 99.62 |
| | Albacore | 0 | 26.64 | 707.10 | 3.8 | 99.97 | 99.25 |
| Staphylococcus Aureus | MSRCall | 0 | **8.17** | **83.97** | 2.9 | 100.00 | **99.97** |
| | SACall | 0 | 9.72 | 449.61 | 2.9 | 100.00 | 99.54 |
| | DeepNano-blitz | 1 | 12.75 | 214.43 | 2.9 | 100.00 | 99.89 |
| | Guppy-KP | 0 | 12.09 | 774.65 | 2.9 | 100.00 | 99.21 |
| | Guppy | 0 | 9.13 | 309.36 | 2.9 | 100.00 | 99.68 |
| | Albacore | 0 | 12.20 | 506.36 | 2.9 | 100.00 | 99.50 |
